# EnhancerNet: a model for enhancer selection in dense regulatory networks recapitulates the dynamics of cell type acquisition

**DOI:** 10.1101/2024.02.03.578744

**Authors:** Omer Karin

**Affiliations:** Department of Mathematics, Imperial College London, London, SW7 2AZ, UK

**Author notes:** Corresponding author: Omer Karin.

## Abstract

Understanding how cell identity is encoded in the genome and acquired during differentiation is a central challenge in cell biology. We derive a theoretical framework called “EnhancerNet” that models dense feedback networks involving transcription factors and enhancers, which can be parameterized from terminal cell identities without fitting unobserved variables. EnhancerNet recapitulates the dynamics of enhancer selection and cell type specification via two distinct pathways: direct reprogramming or differentiation through transient, multipotent progenitor states. These pathways capture the hallmarks of their respective counterparts in animal cells, with the model reproducing known reprogramming recipes and the complex hematopoietic differentiation hierarchy. Using EnhancerNet, we show that hierarchical progenitors emerge as transient states during differentiation and propose a method to predict their identity from terminal states. The model explains how new cell types could evolve and highlights the functional importance of distal regulatory elements with dynamic chromatin in multicellular evolution.

## Introduction

A single fertilised animal egg cell divides and differentiates into many cell types that are maintained throughout the life of the organism. Each cell type is associated with a distinct gene expression profile, which can be acquired by gradual differentiation through progenitor states, or by direct reprogramming (transdifferentiation). Despite great advancements in the molecular characterization of cell dynamics, many fundamental questions regarding cell type acquisition remain open. Specifically, key open challenges include deciphering how cell types and their associated expression profiles are encoded in the genome, identifying the regulatory mechanisms underlying stepwise differentiation versus direct reprogramming pathways, understanding how inputs trigger the acquisition of particular cellular phenotypes, and determining how new cell types with unique expression signatures can evolve.

Experimental evidence suggests that to address these questions, we need to understand the interactions between *transcription factors* (TFs) and cis-regulatory elements known as *enhancers*. TFs are DNA-binding proteins that can modulate the expression of other genes. Specific cell types are associated with the expression of distinct combinations of TFs which play a crucial role in maintaining and modulating cell identity [1–12]. TFs bind enhancers to activate cell-type specific expression patterns. Each enhancer can be bound by many TFs, and, in turn, initiate gene expression in one or more distal target genes [13, 14]. The binding of TFs to an enhancer can change its activity by modulating epigenetic properties such as the biochemical properties of its associated chromatin [15–19]. Empirically, it has been observed that TFs that control cell identity bind enhancers that control their own expression and the expression of other identity-determining TFs that are coexpressed in specific cell types, forming dense autoregulatory networks [4, 5, 7, 20, 21]. These dense autoregulatory networks are likely to play a crucial role in cell type specification, as their enhancers show activation patterns that are highly cell type-specific [4, 22]. This contrasts with the activity of TFs, which may be shared across multiple cell types [4].

While the selection of cell type-specific enhancers is central to cell type acquisition and may have been a major evolutionary innovation underlying multicellularity [23, 24], it is not captured by existing models for cell type acquisition. Rather, a greater emphasis has been placed on TF interactions, focusing on how small transcriptional network motifs can generate cell fate bifurcations [25–29], or how attractor states can emerge in large transcriptional networks [30–34]. While these studies have been important in dissecting specific observations, including bistability [25], multistability [30, 35], reprogramming [33, 34], and multilineage priming [26], they are limited in their ability to make predictions across a broad range of experimental observations, and they do not inform of the role of enhancers with dynamic chromatin for cell fate regulation.

Here we address this by developing an explicit theoretical framework for dense regulatory networks (“EnhancerNet”), incorporating a wide range of experimental evidence on the regulation of the process. We demonstrate that the dynamics of the core regulatory network that controls cell identity can be inferred from the identity of the terminal cell states, propose predictive models for reprogramming and differentiation, and provide a theory for the role of distal regulatory elements with dynamic chromatin for multicellular evolution.

## Results

### Model for enhancer activation dynamics in transcriptional feedback networks

We modeled transcriptional activation by considering a two-step process where (1) transcription is initiated at enhancer *i* by the binding of rate-limiting transcriptional initiation machinery at rate *p*_*i*_, and (2) transcription initiation at the enhancer results in the transcription of gene *j* by a coupling rate *q*_*i,j*_.

For the first step, we assumed that enhancers compete for the binding of transcriptional machinery such as Mediator complex, which can be bound to each of the enhancers (Figure 1A). We denote by *ϵ*_*i*_ the energy for transcriptional activation at enhancer *i*, and assume, for simplicity, that the transcriptional machinery is always bound to one of the enhancers. The equilibrium rate of transcription initiation at enhancer *i* is proportional to the Boltzmann distribution:

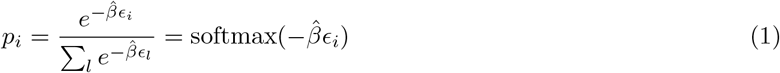

where 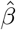 is inverse temperature. The initiation of transcription at enhancer *i* can then result in the transcription of several associated genes, and each gene can be transcribed following an interaction with one of several enhancers (Figure 1A). The dynamics of the expression of gene *j*, denoted *x*_*j*_, is given by the sum:

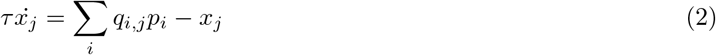

where *τ* is the timescale of gene expression changes. Feedback in the model occurs when TFs modulate the activation energies of the enhancers (Figure 1D). In classical models for gene regulation in prokaryotes, the binding of activator or repressor molecules directly modulates the rate of transcription initiation, corresponding to the variables *ϵ*_*i*_ in our model [36]. Here, we consider the possibility that the dominant mode of enhancer activation is *indirect*, through the modulation of enhancer chromatin. Binding of TFs to enhancers leads to loosening of nucleosomes and to the biochemical modification of histone proteins [15–17], resulting in dynamic changes in enhancer chromatin that are closely linked to enhancer activity [18, 19]. From a biophysical perspective, this can be captured by a model in which *ϵ*_*i*_ is set by a latent variable *m*_*i*_, which accumulates proportionally to the binding of TFs:

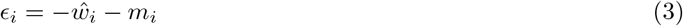

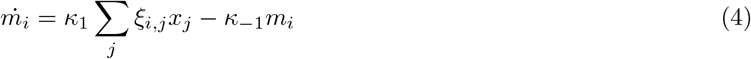

where *ξ*_*i,j*_ is the effective binding rate TF *j* to enhancer *i*, and where *κ*_1_, *κ*_−1_ are the associated “on” and “off” rates for *m*_*i*_. The weight parameter *ŵ*_*i*_ determines the energy in the absence of *m*_*i*_. The mechanism described in Eq. 4 is generic and may correspond to several underlying biological processes and is, in essence, similar to modulation of receptor activation energy by methylation in bacterial chemotaxis [37]. Taking a quasi-steady-state of *m*_*i*_ and denoting by 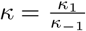, we have the following dynamics (Figure 1B):

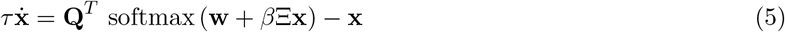

where Ξ is a matrix whose entries are *ξ*_*i,j*_, **Q** is the association matrix whose entries are *q*_*i,j*_, **w** is the bias vector whose entries are 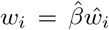, the effective inverse temperature is 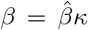, and **x** is the vector of TF expression whose entries are *x*_*i*_. In this study, we will refer to the network model and the associated dynamics as *EnhancerNet*.

**Figure 1:**
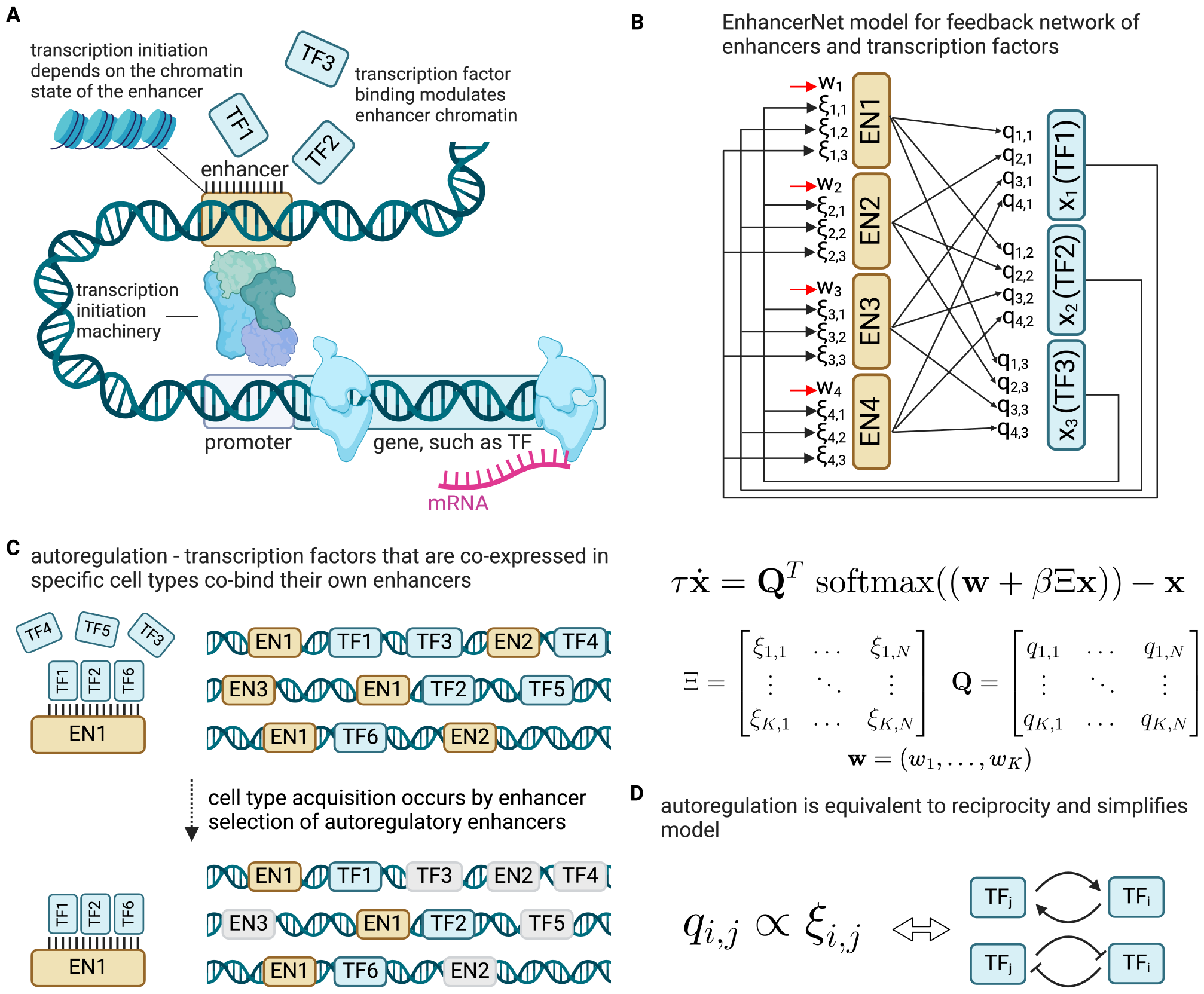
EnhancerNet model. (A) Model setup. Enhancers are cis-regulatory elements that can initiate transcription in distant genes through interaction with specific TFs and transcriptional machinery. Each enhancer can interact with several genes, and each gene can be controlled by multiple enhancers. Transcription initiation depends on the energy required for the transcriptional machinery to bind and initiate transcription from the enhancer. Binding of TFs modulates enhancer chromatin and can increase the transcription rate of the enhancer. (B) EnhancerNet model. Enhancer types (left) bind specific combinations of TFs (right) according to binding strengths specified by the Ξ matrix, and, in turn, initiate transcription according to the rate matrix **Q**, and there is also a weight vector **w**. (C) Autoregulation is the observation that TFs that are co-expressed in specific cell types co-bind their own enhancers. These enhancers are, in turn, selected and activated in these specific cell types. (D) Autoregulation constrains **Q** ∝ Ξ and implies reciprocity.

While Eq. 5 considered different physical enhancers as separate entities, in practice enhancers with similar binding profiles can appear throughout the genome [7, 38], corresponding to similar rows of Ξ. We denote these as enhancer *types* and note that Eq. 5 can be rewritten so that Ξ is a matrix of enhancer types (i.e., with distinct rows) and the matrix **Q** corresponds to the overall association between enhancer types and gene expression (Methods). Hereafter we will assume that Eq. 5 captures the dynamics of enhancer types, which can correspond to the activation of multiple physical enhancers.

Eq. 5 can be further constrained by taking into account well-established properties of developmental networks. TFs associated with specific cell identities form densely interconnected networks in which they co-specific enhancers associated with the cell identity, which in turn activate the same TFs [4, 5, 7, 39] (Figure 1C). In the model, autoregulation implies a strong positive correlation between the rows of Ξ, corresponding to binding profiles to enhancers, and the rows of **Q**, corresponding to the effect of these enhancers on TFs. We capture this by taking **Q** = Ξ (Figure 1D, relaxing this to assume that **Q**, Ξ are merely correlated does not affect our conclusions). We therefore have the following model for the dynamics:

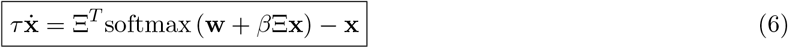

The system described by Eq. 6 is a gradient system, whose dynamics track the gradient of a scalar potential [40] :

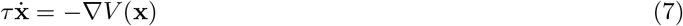

with:

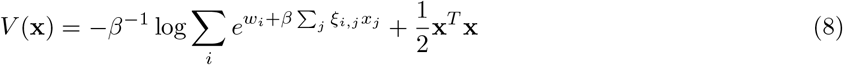

The dynamics captured by Eq. 6 are mathematically similar to models for efficient memory storage in artificial neural networks (Methods) [40], and we will exploit this relationship when we analyze different properties of the dynamics.

### Autoregulation is equivalent to reciprocity in the EnhancerNet model

Autoregulation, captured by **Q** = Ξ, implies that the interactions between TFs in the developmental network are *reciprocal*, that is, for each pair of TFs *k, j*, we have that 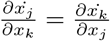 Moreover, reciprocity and autoregulation are effectively equivalent (Methods) (Figure 1D). This equivalence provides a functional explanation to why reciprocal interactions are ubiquitous motifs of developmental networks [26, 27, 41, 42].

Reciprocal interactions can be positive (mutual activation) or negative (mutual repression), and both possibilities are common in developmental networks. In the EnhancerNet model, as well, both possibilities occur. However, there is no direct repression in the EnhancerNet model. In the model, enhancers only enhance transcription, and the binding of TFs to enhancers increases their activity. Repression is thus a byproduct of global inhibition by competition over shared transcriptional machinery.

### Robust specification of combinatorial cell types through interaction between TFs and enhancers

The EnhancerNet model can explain how cell-type specification and enhancer selection occur in the gene regulatory network. At large *β* the dynamics given by Eq. 6 stabilize the expression profiles associated with enhancers, given by the rows of Ξ (or, in the general case, the rows of **Q**, Methods). That is, the stable fixed points of the network are given by the vectors (*x*_1_, …, *x*_*N*_) = (*ξ*_*i*,1_, …, *ξ*_*i,N*_) for all enhancer types (Figure 2A,B). Each stable fixed point corresponds to a cell type with a unique combinatorial transcription factor expression pattern. At these fixed points, cell type-specific enhancers show maximal activity. Thus, enhancer activity is highly cell-type specific, while lineage-determining TFs may be shared between cell types, recapitulating experimental knowledge on the cell-type specificity of enhancers and the cell-type specific activation of autoregulatory transcriptional factor-enhancer networks [4].

**Figure 2:**
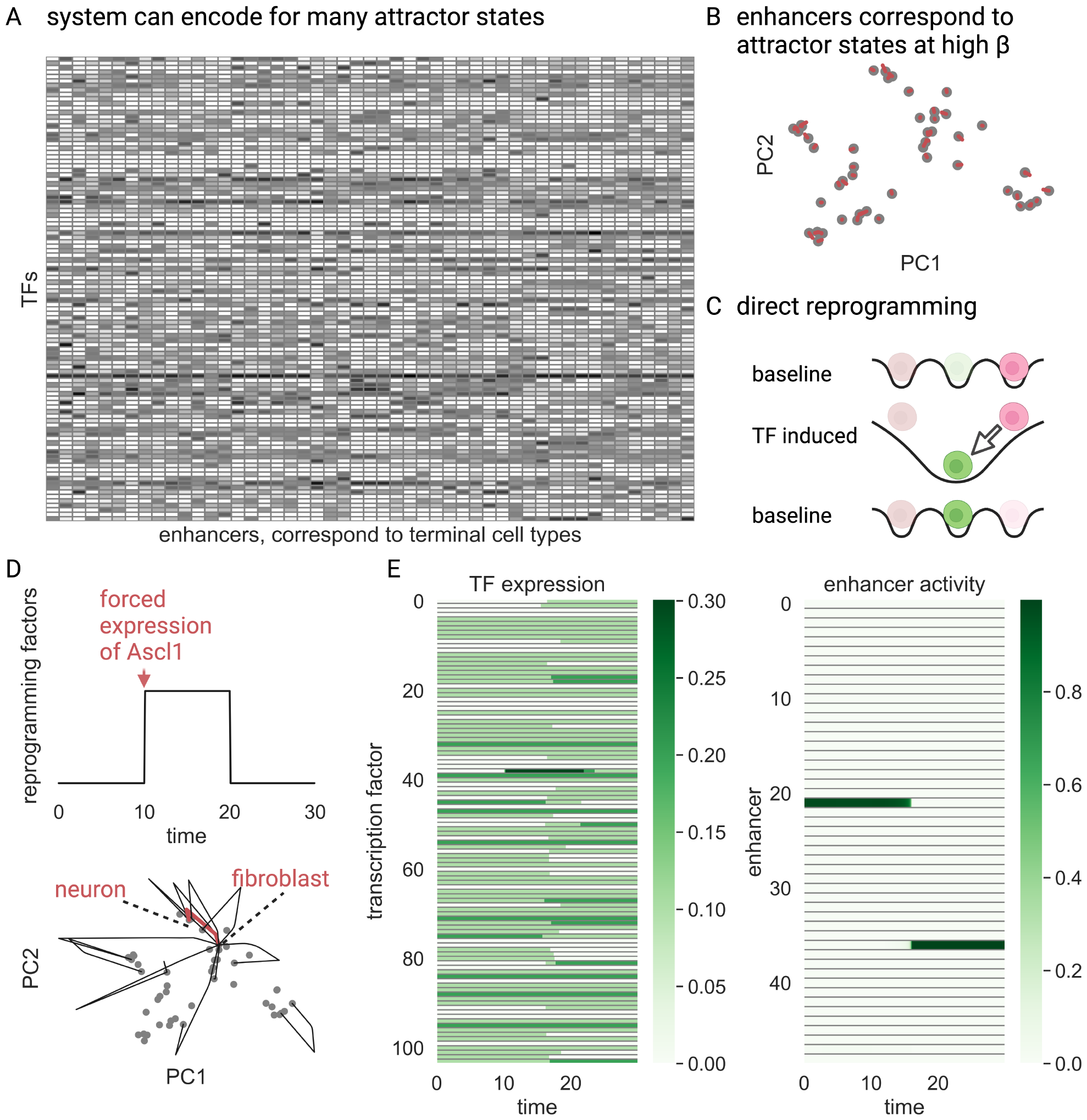
Specification of cell types and direct reprogramming in the EnhancerNet model. (A) The EnhancerNet model was initialized with the TF expression profiles of 49 cell types from Tabula Muris[43], taking 104 TFs with high variability between cell types (Methods). Plotted is the resulting Ξ matrix, with higher values corresponding to darker colors. (B) At high *β* each is row of Ξ an attractor of the network. The panel depicts the PCA transformation of Ξ, with individual rows plotted in gray. They are all attractor states - trajectories that start at the vicinity of each of these states converge to them. Red lines correspond to trajectories starting from rows of Ξ with 25% multiplicative noise. (C) Constitutive overexpression of TFs alters the energy landscape and can result in a bifurcation where cell types lose stability and transition to another cell type. (D) The overexpression of Ascl1 transitions fibroblasts to neurons in the model, capturing the known effect of Ascl1 overexpression in fibroblasts. The reprogramming path is plotted in red (lower panel), with black paths capturing other known reprogramming recipes (Methods). (E) Expression of TFs (right panel) and enhancer activity (left panel), corresponding to the probability that transcription is initiated from the enhancer.

The EnhancerNet model can also explain how the gene regulatory supports the induction of specific cell fates or the restriction of inducible cell-fates. This is predicted to occur through the modulation of the weight vector **w** (Methods). The vector **w** can be controlled, for example, by environmental signals and by the receptors and signaling pathways expressed by the cell, and may be under feedback control. Importantly, changing **w** does not change the identity of the attractor states. Thus, the EnhancerNet model provides a mechanism for specifying both terminal cell types and controlling which cell types can be specified, without interfering with the cell identities.

The observation that cell types correspond to rows of Ξ suggests an intriguing possibility: that Eq. 6 can be parameterized using the values of the observed (terminal) cell types. The weight vector **w** can be parameterized either with identical entries (with all cell fates accessible) or with larger entries corresponding to the accessible cell types. This leaves a single free parameter *β* for the entire regulatory network. The model can thus, in principle, predict the steady-state and dynamics of the entire transcriptional network by only considering the terminal cell fates. In the rest of the manuscript, we will scrutinize this possibility and show that we can indeed reconstruct the complex dynamics of the regulatory network by considering only the terminal cell states.

### Differentiation and reprogramming processes are captured by the model

To test the possibility that the EnhancerNet model can predict the dynamics of the transcriptional network that controls cell identity, we will focus on the processes of cell identity acquisition. These processes can be divided into two broad classes: direct reprogramming, in which there are direct transitions between differentiated cell types, and differentiation processes that start from (or transition through) progenitor states, which may be multipotent.

Direct reprogramming is experimentally carried out by artificially overexpressing TFs, and also possibly non-coding factors or small molecules [11]. Direct reprogramming has been demonstrated between dozens of cell types by overexpressing a wide range of TFs and TF combinations. Two clear distinguishing features for whether specific TFs can efficiently reprogram into a target cell type are that these factors are (a) highly expressed in the target cell type, and (b) their expression is unique to that cell type [44].

The very existence of direct reprogramming is beyond the scope of classical models for cellular differentiation that are based on competition between lineage-determining factors. However, it is a straightforward consequence of the dynamics of Eq. 6 (Figure 2C). The overexpression of a TF *x*_*j*_ modulates the energy of each stored pattern (Methods). A large decrease in the energy of a pattern can increase its basin of attraction, which can then encompass other (previously stable) expression patterns. If a cell has initially been placed in one of these distant patterns, that is, **x**_*i*_ = (*ξ*_*i*,1_, …, *ξ*_*i,N*_), then, after TF overexpression, it can transition directly to a new pattern **x**_*k*_ = (*ξ*_*k*,1_, …, *ξ*_*k,N*_).

Which transcription factor can be most efficient in reprogramming to a specific cell type? The overexpression of the factor *x*_*j*_ by a degree *δ*_*j*_ decreases the energy of pattern *k* by *δ*_*j*_*ξ*_*k,j*_ (Methods). The optimal transcription factor for reprogramming to a target pattern *k* will uniquely decrease the energy of this pattern, while avoiding decreasing the energy of other patterns. Therefore, it is the one for which *ξ*_*k,j*_ is large, corresponding to high expression in the target pattern, and for which *ξ*_*i,j*_ is small for all *i* ≠ *k*, corresponding to low expression in other types of cells. The requirement that a reprogramming factor is unique and highly expressed in the target cell type captures the known properties of reprogramming factors and is the basis of computational methods to identify reprogramming factors [44, 45]. Thus, our model is consistent with direct reprogramming and its known phenomenology.

To test whether the model can recapitulate known reprogramming recipes, we generated a data set for 49 terminal cell identities using Tabula Muris [43] (Methods, Figure 2A). We then set Ξ according to the TFs used in 12 established reprogramming recipes, as well as other TFs with high expression and variability. The transient overexpression of the reprogramming factors recapitulated the known reprogramming behavior, with the system transitioning from the original attractor state (e.g. fibroblast) to an end attractor state in line with experiment (Figure 2D,E). Thus, the EnhancerNet model can quantitatively recapitulate direct reprogramming, with no fitting parameters and by only considering the terminal states.

As a specific demonstration of direct reprogramming, consider the reprogramming of a fibroblast into a neuron by overexpression of Ascl1 (Figure 2E) [46]. The cell begins in a fibroblast state where a specific enhancer type is strongly activated. The transient activation of a specific TF can destabilise this state, resulting in a transition to a neuron state that is associated with the activation of a neuron-specific enhancer type.

### Differentiation through Waddington landscapes with multilineage priming captured by the model and progenitors can be predicted from the identities of terminal cells

Direct reprogramming is an important experimental phenomenon and may also occur in natural settings following tissue perturbation [47]. In homeostatic and developmental settings, however, it is more typical for differentiation trajectories to occur through a series of multipotent progenitor states [48]. Such differentiation trajectories have been extensively studied in mammalian hematopoiesis, where a population of multipotent progenitors can give rise to many blood lineages [49, 50]. Other well-studied examples include gut homeostasis [51], stomach [52], skin [53], as well as throughout development [54].

Differentiation trajectories share several common characteristics. Differentiation proceeds in a directed manner through a series of progenitor states. Each progenitor may acquire one of several terminal fates, with the range of target cell fates becoming more restricted as differentiation proceeds. For example, the cell may initially be in a multipotent progenitor state, and then transition to a bipotent state followed by a unipotent state. These dynamics are most famously conveyed by the Waddington landscape [55], with the image of a ball rolling down a hill segregated by valleys capturing the progressive restriction of cell fate as it progresses through transitional progenitor states. Progenitors co-express at low levels the lineage-determining TFs associated with their target fates, a phenomenon known as multilineage priming [51, 54, 56–60]. Although the Waddington landscape image has gained great popularity for conceptualizing the dynamics of cell fate specification and for developing quantitative models for differentiation dynamics [55, 61, 62], it is not clear how these complex hierarchical dynamics are implemented by the gene regulatory network.

Here we show that Waddingtonian cell fate specification dynamics and the existence and identity of the progenitor states is an emergent property of the dynamics captured by the EnhancerModel. They are due to the second mechanism for cell type transitions in the model - *annealing*, where *β* is transiently decreased by the cell (“heating up”) and then slowly increased (“cooling down”) (Figure 3A). Recall that *β* is a course-grained parameter that depends on global properties of chromatin accessibility and can be modulated by global changes in mechanisms that affect the latent modification *m*. Decreasing *β* results in a widespread pattern of chromatin accessibility and gene expression, while increasing *β* results in the activation of more specific enhancers, in line with experimental knowledge of differentiation hierarchies [63]. Specifically, decreasing *β* destabilises terminal attractor states, while increasing *β* restabilises them, resulting in a transition towards a new terminal state (Figure 3).

**Figure 3:**
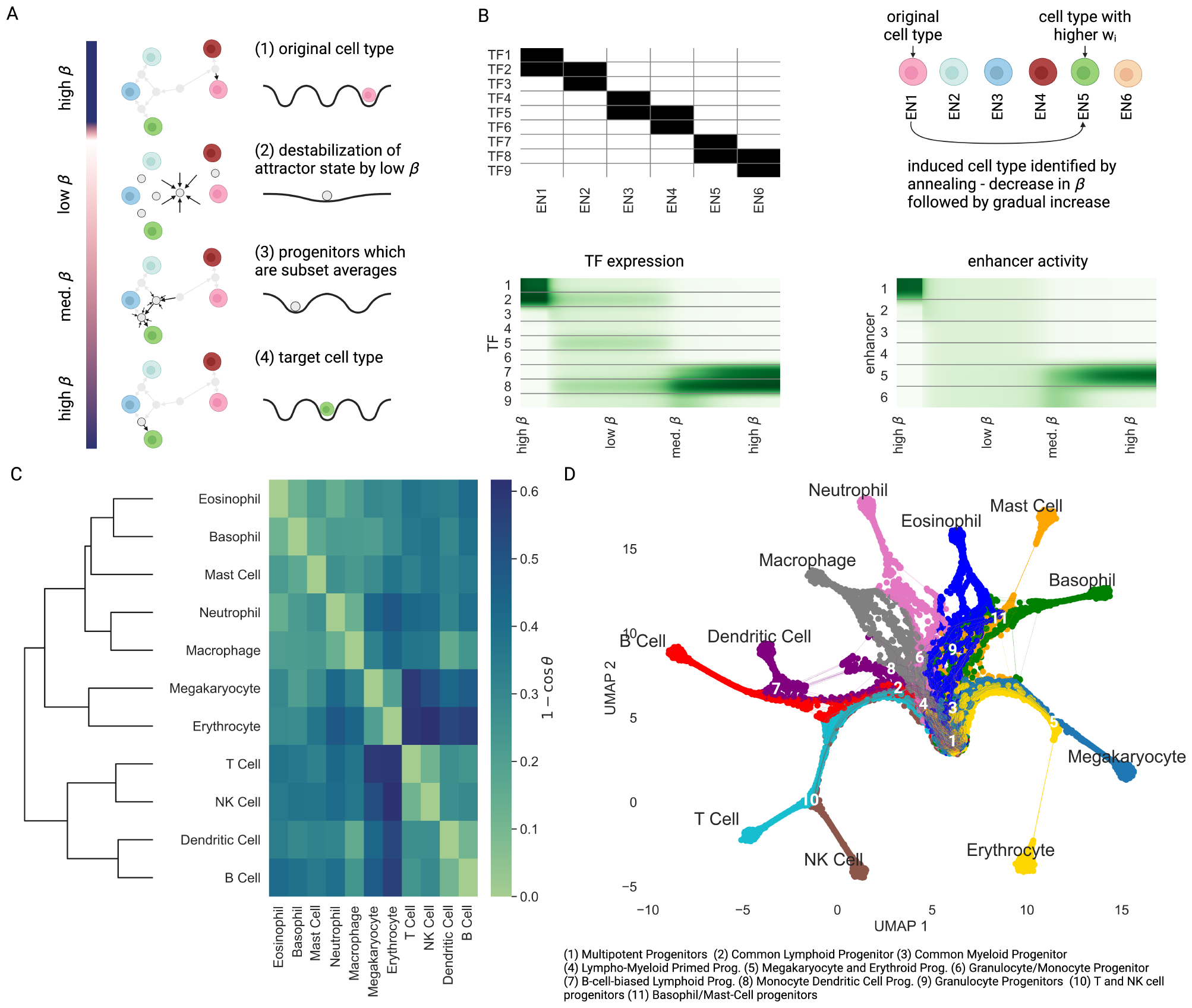
Hierarchial differentiation by annealing. (A) Annealing is the process by which *β* is transiently decreased and then increased. The decrease in *β* transitions the cell to a global attractor state corresponding to a multipotent progenitor with an averaged expression profile of terminal cell types. As *β* increases again, new attractor states are created, which are averages of smaller subsets of the terminal cell types, corresponding to progenitor states with limited potency, until finally, the cell transitions to a terminal cell type. (B) Progenitor states appear in subsets of cell types that have high cosine similarity. In this example, we considered a simple setup where enhancers bind overlapping TFs (upper left subpanel, with positive TF binding denoted in black), the dynamics begin from a cell type corresponding to the binding profile of En1, and En5 has a slight positive weight *w*_5_ = 0.05. Annealing was simulated as in the middle right subpanel. The dynamics through progenitors that show multilineage priming (lower left subpanel) and a transition from widespread enhancer activity to cell type-specific activity (lower right subpanel). (C) The cosine similarity of the expression profiles of the terminal blood lineages, where expression profiles were calculated using the TFs that show both sufficiently high interlineage variability (Methods). Hierarchial clustering according to cosine similarity recreates the “classical” hematopoiesis differentiation tree. (D) UMAP plot of differentiation trajectories, generated by direct simulation of annealing in an EnhancerNet model calibrated by the hematopoietic terminal lineage expression profiles. Trajectories were simulated from an initial homogeneous population with the addition of noise and with **w** adjusted to produce a balanced differentiation profile (Methods). Points represent samples of the trajectories at constant time intervals, with the color corresponding to the identity of the final state. Simulation recreates the major observed progenitor states in hematopoiesis, including deviations from tree-like differentiation.

The annealing process proceeds as follows. The accessible terminal cell states, given by the rows of Ξ where **w** is sufficiently large, are all stable when *β* is very large, while their global average is the only stable state when *β* is small. This state expresses at low levels the TFs associated with all target lineages. It corresponds to a multi-potent progenitor cell identity. As *β* increases, this state loses stability, and new steady states appear transiently. These new steady-states are averages of increasingly restricted groups of terminal cell states, and they correspond to progenitors of restricted potential. Annealing thus recapitulates the hierarchical Waddingtonian differentiation dynamics and low-level multilineage progenitor expression patterns.

While progenitors generally correspond to averages of terminal cell types, not all averages are equally likely to be observed and stable progenitors. Rather, the stable progenitor states correspond to subsets of terminal states with high internal cosine similarity. As an illustrative example, we considered a simulation of a regulatory network with 9 TFs, indexed *x*_1_, …, *x*_9_, and 6 enhancers, indexed Ξ_1_, … Ξ_6_. Each enhancer can be bound by 2 TFs, and enhancers 1-2, 3-4, and 5-6 overlap by a single TF (Figure 3B). The initial state corresponded to an expression profile associated with Ξ_1_ and a large *β*. We also take a slightly larger weight *w*_5_ for Ξ_5_. Annealing (decrease in *β* followed by gradual increase) transitions the cell through two progenitors - a “multipotent progenitor” (low *β*) which corresponds to the average expression of all the enhancers; a “restricted potential progenitor” (med. *β*) associated with the average expression of Ξ_5_, Ξ_6_, and finally differentiation to Ξ_5_. Thus, differentiation occurs through Waddingtonian dynamics associated with multilineage priming. Annealing also provides an efficient strategy to detect a global energy minimum corresponding to an induced cell type.

The annealing model makes the specific prediction that differentiation hierarchies, including the identity of observed progenitor states, can be estimated by only knowing the cosine similarity between the identities of the terminal cell states. The model specifically predicts that they will appear only between transcriptional profiles with high internal cosine similarity. To test this, we considered differentiation in hematopoiesis, a system in which differentiation trajectories are complex and have been extensively studied. Using data on hematopoietic lineages in mice (extracted from Haemopedia [64]), we considered the expression profile of all TFs that showed variability between terminal lineages and used the model to estimate the differentiation hierarchy that generated these lineages.

We tested the hypothesis that progenitors emerge between correlated expression profiles by performing hierarchical clustering according to pairwise cosine similarity between the expression profiles of terminal cell types (Figure 3C). Hierarchical clustering produces a dendrogram (tree) that shows how expression profiles are progressively grouped into clusters based on their cosine similarity. Hierarchical clustering resulted in a tree structure (dendrogram) that recreates the classic hematopoietic hierarchy (Figure 3C). Branching points at the dendrogram capture the Common Lymphoid Progenitor (CLP), Common Myeloid Progenitor (CMP), Megakaryocyte and Erythroid Progenitor (MEP), Granulocyte and Macrophage Progenitor (GMP), T and NK cell progenitors (TNK), and B cell-biased lymphoid progenitor (BLP). Thus, a simple unsupervised clustering method with access to only the terminal cell fate information can reproduce a faithful estimate of the known complex differentiation landscape in hematopoiesis.

Although hierarchical clustering produces a tree-like structure that recapitulates the main progenitor states in hematopoiesis, differentiation trajectories in the model need not be tree-like. As *β* increases, multiple steady states can appear and disappear, and these new steady states may be averages of overlapping terminal states. The key prediction of the model is that these states will correspond to averages of terminal states with high internal cosine similarity. This can explain observed deviations from tree-like dynamics in hematopoiesis. For example, dendritic cells can emerge from both the myeloid lineage through macrophage-dendritic progenitors and from the lymphoid lineage through B cell-biased lymphoid progenitors [65]. This observation is fully in line with our model, as the gene expression profile of dendritic cells has a high cosine similarity to B cells, but also macrophages (and, more generally, myeloid cells), while B cells are less similar to myeloid cells (Figure 3C). Other deviations from a tree structure can be captured similarly by the model. For example, lymphoid and granulocyte cells have moderate cosine similarity, and granulocyte and erythro-megakaryocytes have moderate cosine similarity, but lymphoid and erythro-megakaryocytes are poorly correlated. Thus, each of the first two might share accessible progenitors, while the latter will be less likely to. The two progenitors indeed exist (Lympho-Myeloid Primed Progenitors and Common Myeloid Progenitors) while the latter, to the best of our knowledge, does not. Thus, the EnhancerNet model can quantitatively explain deviations from tree-like differentiation patterns in hematopoiesis.

To test whether EnhancerNet can recapitulate differentiation trajectories in hematopoiesis, we simulated annealing trajectories starting from a homogenous initial population, corresponding to hematopoietic stem cells. To capture the balanced production of multiple cell types in hematopoiesis, we added noise to Eq. 6, and implemented a negative feedback routine to tune the weight vector **w** (Methods). This resulted in dynamics that generated all cell fates in a balanced manner. We then plotted the differentiation trajectories on a UMAP plot (Figure 3D), revealing that differentiation dynamics proceed through a series of multipotent progenitors corresponding to the progenitors observed in hematopoiesis. Thus, the model quantitively recapitulates differentiation in hematopoiesis with no fitting parameters.

### Evolution of new cell types

We conclude this study by considering the question of *genesis*, namely, how can new cell types, corresponding to attractor states of the EnhancerNet, evolve?

Although in principle a new cell type may arise *de novo*, in practice it is likely to appear by the diversification of an ancestral cell type into sister cell types, a process known as genetic individuation [10, 66]. In this process, a stable attractor state associated with a specific combination of TFs evolves into new and distinct coexisting attractor states associated with new TF combinations.

The EnhancerNet model has unique properties that support the process of genetic individuation of cell types; we propose that these properties may have been instrumental for the evolution of distal cis-regulatory elements with dynamic chromatin in animals. These properties are (i) the ability to support multiple coexisting, highly correlated attractor states, and (ii) the ability to have multiple enhancers regulating the same genes.

The importance of the first property is clear when we consider that around the time of the initial diversification of the ancestral cell types, the sister cells have highly correlated expression patterns. At this stage, they need to coexist as distinct attractor states to be expressed in the body. Thus, in the biological context, the ability to support multiple correlated attractor cell type states is crucial.

The second property allows the regulatory network to generate and modify specific cell types without interfering with other cell types that may have common TFs. This relates to the modularity of encoding cell types by enhancer sequences. We illustrate this by a simple example where an ancestral cell type expressing TFs 1,2,3,4 is diversified into two cell types that express 1,2,3 or 1,2,4 (Figure 4A,B). The initial attractor state is associated with an enhancer type, denoted EN1, that binds all four TFs. In our example, this enhancer is initially placed in the genome near all four TFs. We also assume that multiple copies of EN1 are near TF1 and TF2 (Figure 4C). This phenomena, known as *shadow enhancers* [38, 67, 68], is widespread and well-established experimentally, and its observed behavior is in line with model predictions (Methods). This setup is illustrated in Figure 4C, and corresponds to a stable attractor where TF1-4 are active.

**Figure 4:**
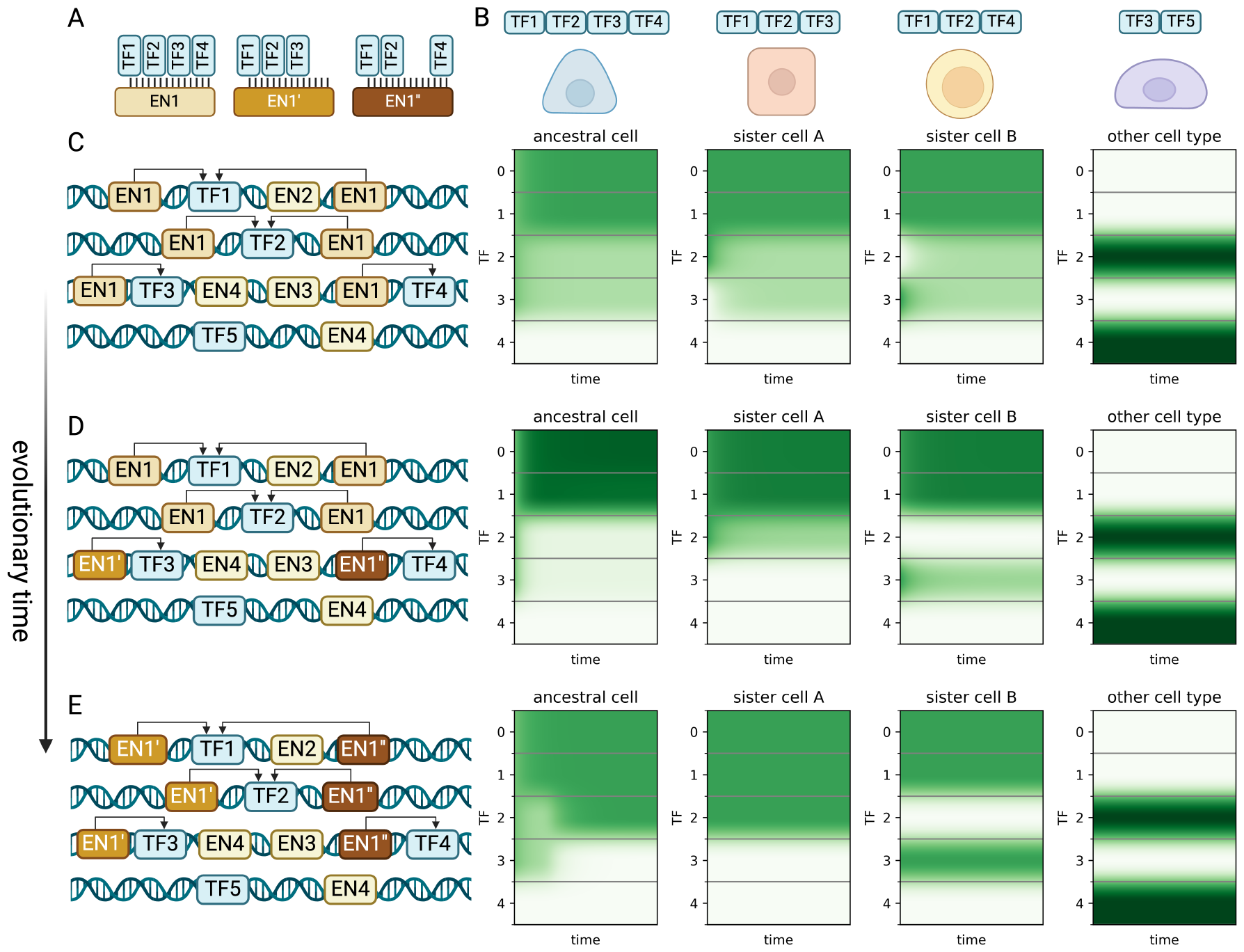
Evolution of new cell types in the EnhancerNet framework. (A,B) We consider the evolution from a cell type associated with the binding pattern of TF1-4 (EN1 in panel A, left cell in panel B) to two sister cell types, each associated with the binding of TF1-2 and either TF3 or TF4 (EN1’ and EN1” in panel A, middle cells in panel B). This must occur without interference with other cells in the regulatory network that may share TFs with these cell types. (C,D,E) The evolution occurs in the presence of other cell types in the gene regulatory network. (C) The original setup (left panel) stabilises only the ancestral cell type - see right panel with simulations starting from the ancestral state where TF1-4 are active, from *specialized A* where TF1-3 are active, and from *specialized B* where TF1-2 and TF4 are active. (D) Mutations in specific enhancers can stabilise states near all cell types. (E) Further mutation can destabilise the ancestral state. Note that for all simulations we used the full dynamics of Eq. 5, with the shade of green corresponding to expression strength of the TF.

Let us consider the possibility that enhancers near TF3 and TF4 evolved, so that the enhancer near TF3 does not bind TF4, and the enhancer near TF4 does not bind TF3 (Figure 2D). Now, there are three stable states - one where TF1-4 are active (with reduced activity of TF3,4) and two where TF1-2 and either TF3 or TF4 are active, corresponding to the new sister cells. Over time, the shadow enhancers may evolve further, leading to the destabilisation of the original attractor state (Figure 2E). Throughout this duration, the other cell types remain stable attractors. Thus, the EnhancerNet model allows the evolution of new cell types without interfering with existing cell types.

## Discussion

Here we derived EnhancerNet - a minimal predictive model for the dynamics of the entire transcriptional network for self-type regulation, based on interactions between TFs and enhancers. The model incorporated several features of the architecture of the regulatory network, namely, that enhancers form dense autoregulatory networks with TFs and that enhancer activity is determined by its chromatin state, which is set by TF binding. These features are sufficient to derive a simple and tractable model that can be used without the need to fit unknown parameters, and that recapitulates the known processes of cell type specification, acquisition, and differentiation.

The mechanism underlying enhancer selection in EnhancerNet is mathematically related to a recent model for memory storage and retrieval known as Dense Associate Memory (DAM), or Modern Continuous Hopfield Networks [40, 69]. Hopfield networks are well-established models for dynamical systems that encode combinatorial attractor states (patterns) through the interactions of their components [70–72]. In the case of transcriptional networks, the attractor states are combinations of TFs. Classical Hopfield networks are based on direct and additive interactions between components [72]. They have limited storage capacity (sublinear in the number of system variables) and cannot store correlated expression patterns. Classical Hopfield networks have long been considered important conceptual models for memory storage and retrieval in the brain [69, 72], and, more recently, they have been employed as conceptual models for cell fate specification, and as the basis of computational methods to study cell state dynamics [33, 34, 73, 74]. It was not clear how such a network would be implemented mechanistically in cells. DAM, in contrast with classical Hopfield models, can store many correlated patterns through higher-order interactions [69, 75]. The mathematical analogy between DAM and enhancer-TF interactions in EnhancerNet provides a mechanism for how cells can retrieve many attractor patterns encoded by a single regulatory network.

The model makes specific testable predictions for processes of reprogramming and differentiation, based on the identities of the terminal states. The model can predict reprogramming ‘recipes’ - which TFs can transition a cell from a given cell type to a target cell type. These predictions are consistent with known algorithms for this purpose and recapitulate established reprogramming recipes. For differentiation, the model predicts the identity of progenitors and can predict complex differentiation hierarchies by using only the identities of the terminal cell types, recapitulating knowledge on differentiation in hematopoiesis. The model also offers a simple heuristic to predict differentiation hierarchies that is based on cosine similarity between terminal expression profiles. To obtain the predictions, we used only minimal preprocessing by considering all variable TFs, using the mature cell identities, suggesting that the approach is robust and does not require a-priori knowledge of binding patterns of TFs. It can also be used for prediction when only limited data is available. EnhancerNet can thus provide a basis for computational methods for predicting differentiation and reprogramming dynamics. The inverse temperature parameter *β* plays a key role in specifying the cell types in the model. At high *β*, cell types associated with enhancer binding patterns are stabilised, while at low *β*, a global average corresponding to a multipotent progenitor is the only stable state. At intermediate *β* averages of smaller subsets of cell types may be stabilised, corresponding to restricted progenitors. The annealing strategy, where cells transiently decrease *β* and then gradually increase it, provides a mechanism for cell type transitions that captures the main hallmarks of hierarchical differentiation and that recapitulates known complex differentiation hierarchies. From a functional point of view, this strategy corresponds to well-established techniques from physics and optimization to settle a system at a global minimum [76, 77]. From a biological perspective, this allows progenitor and stem cells to identify induced cell types (illustrated in Figure 3B) and direct their differentiation towards them, avoiding convergence to metastable states.

Our analysis thus predicts that differentiation from a stem to a terminal cell identity occurs by annealing, that is, by decreasing and then increasing *β*. In our model, *β* is related to the turnover of a latent chromatin modification *m*. Increasing *β* can occur by increasing its global production (*κ*_1_) or decreasing its global removal (*κ*_−1_). The mechanisms underlying the modification of enhancer chromatin involve the activity of many cellular factors, including chromatin remodelers such as the SWI/SNF, Mi-2/NuRD, SET1/MLL, and Polycomb complexes, and involve various mechanisms including the depletion and modification of histone proteins [78–82]. In addition to these, the methylation of DNA itself can play a role in regulating enhancer function [83]. While enhancer activity is tightly linked to specific chromatin signatures associated with TF binding to the enhancer, the underlying mechanisms that cause these signatures and their contribution to enhancer activity are highly complex and not yet fully understood. For this reason, we opted for a minimal model and considered *m* as an (unknown) latent modification, although in practice there are likely to be several underlying factors that contribute to enhancer activity.

How can annealing be implemented in cells? One possibility may be through a positive feedback mechanism, where the number of active genes or enhancers itself decreases *β*. In such a case, the transition from a multipotent progenitor to a more restricted progenitor will increase *β*, pushing *β* higher until the terminal attractor states are reached. Furthermore, *β* can fluctuate during the cell cycle, increasing specifically in G1 where differentiation commitment occurs [84–86]. Thus, cells with a sufficiently long G1 can transition between attractor states, in accordance with the known relation between G1 length and differentiation [85]. While these mechanisms remain putative and need to be experimentally tested, they provide a possible blueprint for ways by which enhancer selection and cell type determination may be carried out.

Finally, the model can explain how new cell types can evolve without affecting pre-existing cell types encoded by the network. This applies both to evolutionary dynamics between generations of organisms and to the evolution of cells in diseases such as cancer. The predictions of the model could be tested by synthetically engineering new cell types. Importantly, the model aligns with the experimentally observed robustness of shadow enhancers, which are thought to play a crucial role in cell type evolution.

In conclusion, we propose that the EnhancerNet model bridges the gap between theoretical and computational models for attractor networks and the rich experimental findings in animal development and physiology, providing a simple predictive framework for modelling whole-cell dynamics of cell fate networks, and explaining the origins of hierarchical differentiation in animal cells.

## Open Access

For the purpose of open access, the author has applied a Creative Commons Attribution (CC BY) licence to any Author Accepted Manuscript version arising from this submission.

## Acknowledgments

We are grateful to Anton Grishchekcin, Benjamin Simons, and Tal Agranov for helpful discussions and for providing feedback on the manuscript.

## Resource and code availability

This paper analyzes existing, publicly available data, with sources cited in the manuscript. The original code for the simulations presented in this study is available from https://github.com/karin-lab/enhancernet. In addition to this code, we have also prepared a suite of computational tools based on the EnhancerNet model, including simulations of reprogramming and differentiation by annealing.

## Methods

### Equivalence of model with physical enhancers to model with enhancer types

We consider the dynamics of Eq. 5 where a subset ℒ of enhancers has identical association patterns, that is, *ξ*_*i,j*_ = *ξ*_*k,j*_ = *ξ*_*j*_ for all *j* and all *i, k* ∈ ℒ. Then:

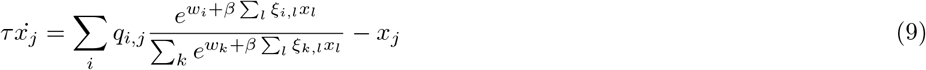

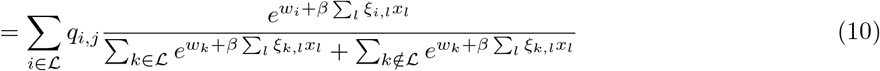

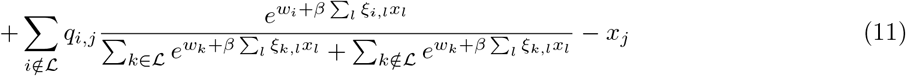

Now:

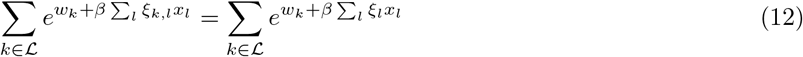

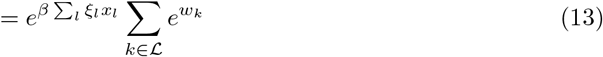

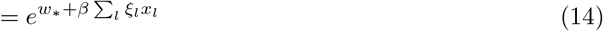

where 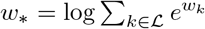 Thus:

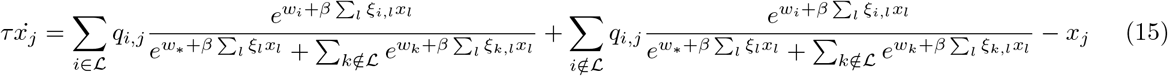

Denoting the denominator by *Z*, we notice that:

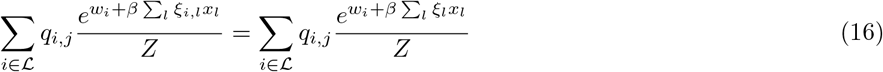

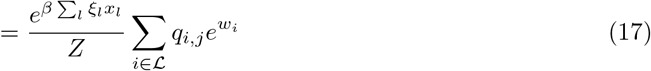

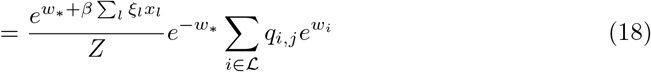

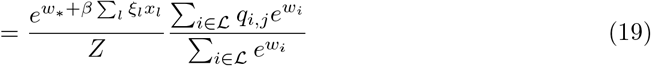

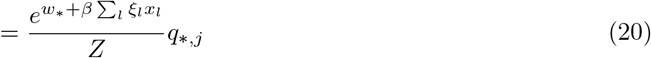

where 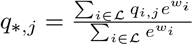 Thus, we have that:

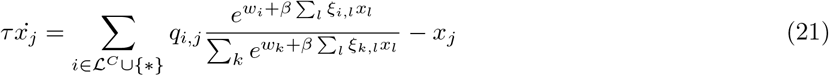

which does not include the physical enhancers of type ℒ, which are replaced with a combined “enhancer type” with the association constant *q*_*∗,j*_ and weight *w*_*∗*_.

### Reciprocity and autoregulation in enhancer feedback networks

Here, we will show that reciprocity:

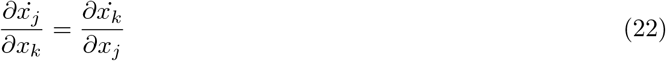

for all *k, j* implies that *q*_*i,j*_ = *ξ*_*i,j*_ for all *i, j*. Since the above relation is trivial for *j* = *k*, we take *j* ≠ *k*:

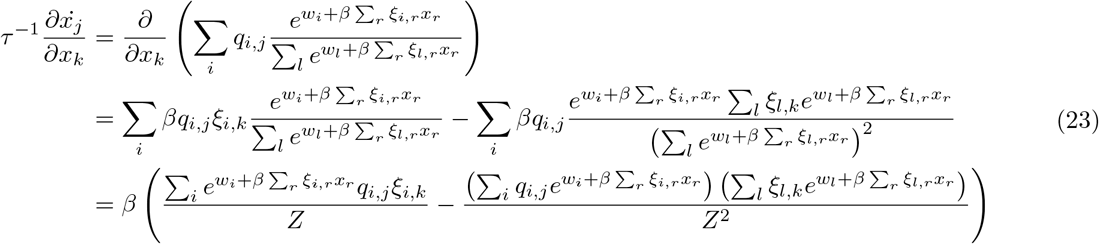

where 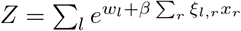 Thus, for an arbitrary *x*, the equality 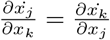 implies that *q*_*i,j*_ = *νξ*_*i,j*_:

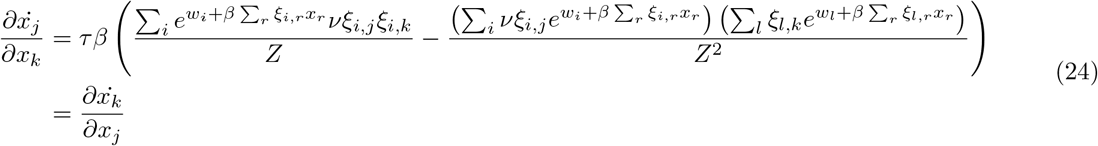

Thus, reciprocal interactions between TFs are equivalent to the autoregulation of TFs by binding to their enhancers.

### Scalar potential for transcriptional dynamics

Many derivations related to Eq. 6 appear in Ramsauer et al. [40], including stability analysis of the fixed points, as well as storage capacity and equivalence with other machine learning models. Here, for completeness, we will derive the important results that pertain to our manuscript, and refer the reader to Ramsauer et al. for more in-depth derivations that pertain to other aspects of the model. Considering the symmetric case, we will show that the dynamics are a gradient flow. A scalar potential for Eq. 6 is:

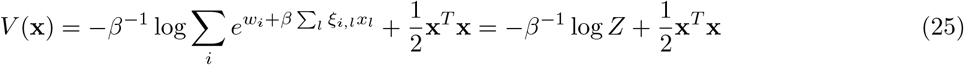

where 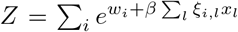 This function is similar to the Lyapunov function of Ramsauer et al. [40], and accounts for the bias **w**. To see that the dynamics are a potential flow:

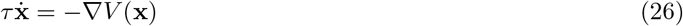

observe that:

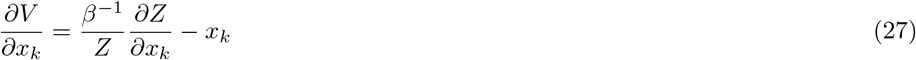

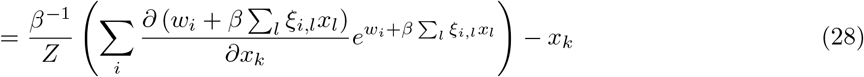

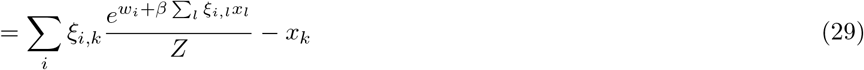

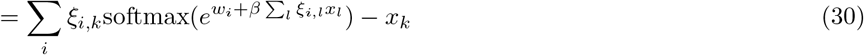

which provides the dynamics of **x**.

### Rows of Q are fixed points at high *β*

Consider the row vectors of Ξ given by Ξ_*i*_ = (*ξ*_*i*,1_, …, *ξ*_*i,N*_). The dynamics of Eq. 6 at **x** = Ξ_*k*_ are given by:

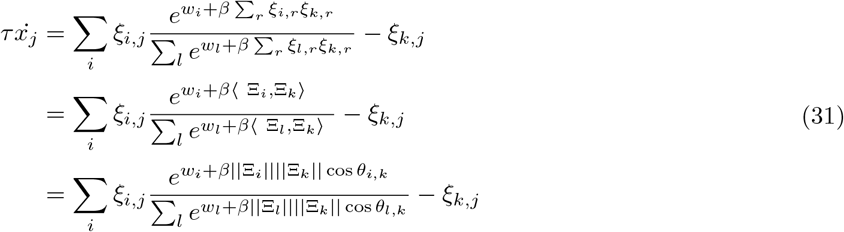

where *θ*_*i,j*_ determines the angle between Ξ_*i*_, Ξ_*j*_ and the magnitude is defined by the Euclidean norm. We assume that the rows of Ξ are all of magnitude unity, which is equivalent to assuming that they have comparable overall binding affinities. Note that this requirement may also correspond to biophysical constraints, such as an overall maximal placement rate of the latent modification *m* on the enahncer. In this case:

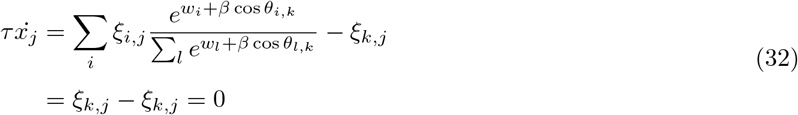

where the last equality holds for sufficiently large *β* and assuming the rows of Ξ are distinct, since in that case cos *θ*_*i,k*_ = 1 if and only if *i* = *k*. Thus, for a large enough *β*, the rows of Ξ are fixed points of the dynamics. What about the case where the matrix **Q** is distinct from Ξ? Consider then dynamics at **x** = **Q**_*k*_ :

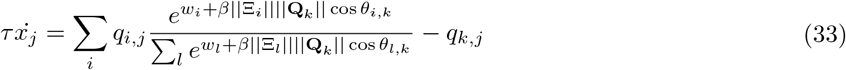

where now *θ*_*i,j*_ determines the angle between Ξ_*i*_, **Q**_*j*_. Thus we have that **Q**_*k*_ is a fixed point of the dynamics at sufficiently large *β* when there is high cosine similarity between Ξ_*k*_, **Q**_*k*_ compared with the other rows of Ξ.

### Averages of rows of Q can be fixed points

Let us denote by ℐ a subset of indices of enhancers of magnitude |*I*| and consider the averaged vector:

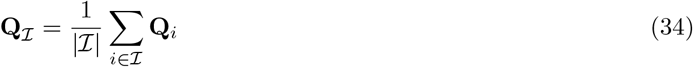

Setting **x** = **Q**_ℐ_, we have the dynamics:

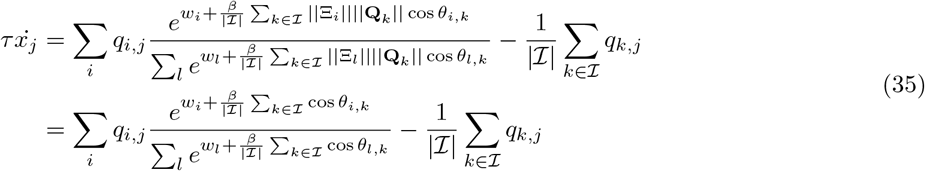

where in the last equality we assumed that the rows of **Q** are also unity. In the case where all weights *w*_*i*_ are similar, this will be zero trivially when *β* is very small and where ℐ is the set of all enhancers (all *K* rows of Ξ), since in this case:

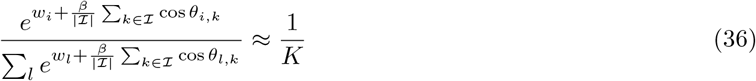

and this can be generalized in a straightforward manner to the case where *some* of the weights of *w* are large and of comparable magnitude. Otherwise, and assuming for simplicity that **w** = 0, we will require that:

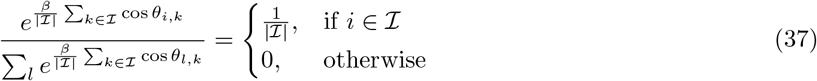

which, at large enough *β*, occurs when the averaged cosine similarity of a set element with the other elements of the set:

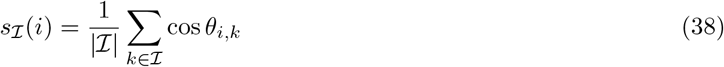

is both (a) comparable within the group, *s*_ℐ_(*i*) ≈ *s*_ℐ_(*r*) for *r, i* ∈ ℐ, and (b) larger than outside the group, *s*_ℐ_(*i*) > *s*_ℐ_(*r*) for *i* ∈ ℐ, *r* ∉ ℐ. An averaged pattern of a subset will therefore be a fixed point if each pattern in the subset is similar to each other pattern, and distinct from patterns outside the subset. Condition (a) is specifically easy to satisfy for subsets with only two patterns; in this case, their average (a “bipotent progenitor”) will be a fixed point when they have large cosine similarity and are distinct from other patterns.

### Global stability of averages depends on *β*

To probe the global stability of averages of rows of **Q**, we can use the scalar potential *V*. The dynamics proceed from high *V* to low *V* so we expect that states with higher *V* are less likely to be stable than states with lower *V*. Note that since the potential function is only defined for the symmetric case, we define the states as averages of the rows of Ξ, which are analogously defined as Ξ_ℐ_ where ℐ is as before a set of indices:

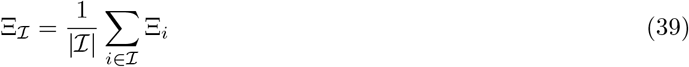

Setting **x** = Ξ_ℐ_, we evaluate:

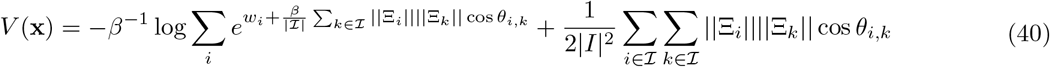

Again taking the magnitude of the rows of Ξ to be unity and **w** = 0 (equiv. to enhancers with similar inding strengths and weights), we have that:

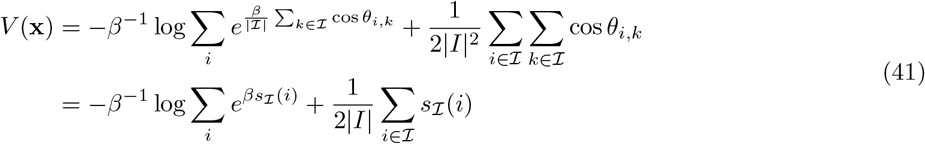

the energy of averaged patterns thus depends only on their cosine similarity relative to all patterns (first term) and internal cosine similarity (second term). For simplicity, let us assume that *s*_ℐ_(*i*) = *A* for *i* ∈ ℐ and *s*_ℐ_(*i*) = *B* for *i* ∉ ℐ (considering that, in general, *A, B* are between (−1, 1)). Then:

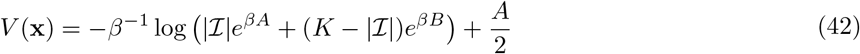

where *K* is the overall number of patterns. For comparable *A* ≈ *B*, corresponding to random subsets, we have that:

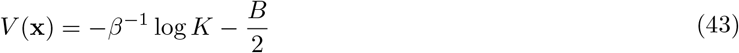

which for large *β* would correspond to 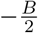 and will thus have high energy when *B* is low. On the other hand, in the case where *A* ≫ *B*, we have that, as a rough approximation:

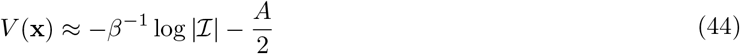

when *β* is very large the size of the subset |*I*| does not contribute to *V* and thus it is minimized when *A* is maximized, that is, at single patterns with *A* = 1 (where *V* = −1*/*2). At intermediate *β*, however, larger subsets may have lower energy. For two subsets where |ℐ_1_| > |ℐ_2_| and *A*_1_ < *A*_2_, we have that the larger group will have lower energy when:

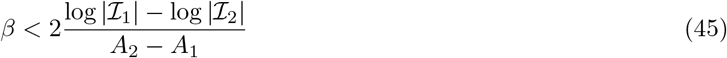

which occurs for a larger range when the difference between the group magnitudes is larger and the difference between their average cosine similarity is smaller. Specifically, the differentiation from a bipotent progenitor |ℐ_1_| = 2 to a terminal identity |ℐ_2_| = 1, *A*_2_ = 1 occurs at a critical *β*:

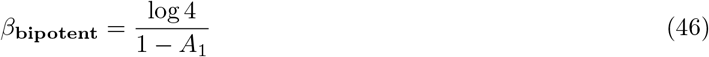

Thus, an annealing strategy where *β* is decreased and their slowly increased results in a transition from large (global) averages, corresponding to multipotent progenitors, to a set of ever more restricted progenitors, similar to the “Waddingtonian” dynamics.

### Direct reprogramming through the over-expression of TFs

Consider a transcription factor *x*_*j*_ that is constitutively overexpressed to an extent *δ*_*j*_. This constitutive expression alters energy levels *ϵ*_*i*_ without feedback. We therefore consider only the effect of the additional TF production on the energy levels of the enhancers, without altering the coordinates of the stable fixed points. We denote by *V* ^*′*^ the potential under the perturbed dynamics. Evaluating *V* ^*′*^ at pattern *k* yields, at high *β*:

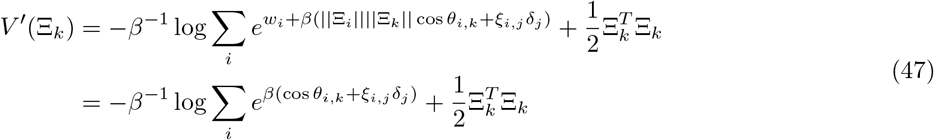

where for simplicity we took **w** = 0, and again assumed that ||Ξ_*i*_|| = 1 for all *i*. Since an individual entry will be of order *ξ*_*i,j*_ ≪ 1, we can assume that for large *β*, the summand takes its maximum at *i* = *k*, where:

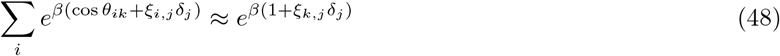

and thus:

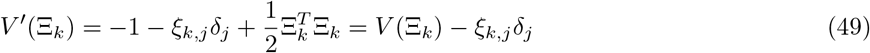

Note that Eq. 49 readily generalizes to a perturbation in several TFs *j* ∈ 𝒥:

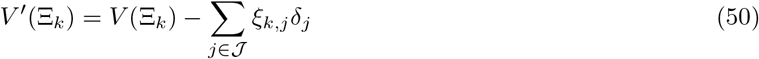

If the goal is to steer the dynamics toward a pattern *k* then the TFs 𝒥 and their overexpression need to be chosen such that 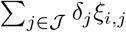 is sufficiently larger for *i* = *k* than for all other patterns *i* ≠ *k*. That is, the ideal expression for a TF *j* is such that *ξ*_*i,j*_ is large for *i* = *k* (amplifying *δ*_*j*_) and zero for *i* ≠ *k*.

### Preprocessing and simulation

The matrix Ξ was initialized from transcript counts extracted from murine cells and averaged over cell type. For the Tabula Muris dataset [43], we used all available data across organs and averaged over cell type. For hematopoiesis, we used the Haemopedia dataset [64] and averaged all cells that belong to the same terminal lineage. We then log-transformed the data (using a log 1 + *x* transformation) and filtered TFs with mean and std expression larger than log 4, as well as for Tabula Muris the TFs that participate in the tested reprogramming pathways (a full list of cell types and TFs used is attached as Supplementary Tables). Although in principle a log transformation is not needed for our model, which does not assume log-transformed values of *x*, it has the statistical advantage of reducing variance in the rows of Ξ and thus reducing sensitivity to TF choice. Finally, the rows of Ξ were normalized to unity. Simulations were performed using Python, taking *τ* = 1, and the (terminal) *β* was chosen so that all patterns were stable.

### Simulation of reprogramming

To simulate forced TF expression during reprogramming we used the following dynamics:

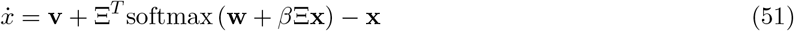

where **v** = {*δ*_1_, …, *δ*_*N*_} is a vector where *δ*_*i*_ corresponds to the degree of activation of TF *i*. In general, we set *δ*_*i*_ = 1 when the TF was in the reprogramming pathway and *δ*_*i*_ = 0 when it was not. However, for a few genes setting *δ*_*i*_ as different from unity was necessary to achieve the correct reprogramming (setting *δ*_*i*_ = 1 achieves reprogramming to a closely related cell type). The full list of reprogramming pathways and relevant *δ* values is attached as a Supplementary Table.

### Simulation of balanced differentiation

Hematopoiesis was simulated by using annealing from *β* = 0 to *β* = 50 over 50 time units. The initial state corresponded to the averaged value of all terminal states. Simulations were performed by adding additive white noise:

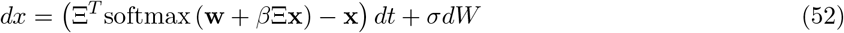

Where we set *σ* = 0.01. To simulate balanced differentiation in hematopoiesis we used the following feedback procedure that aims to imitate actual (in vivo) feedback on differentiation. **w** was initialized as a zero vector. Then, running from *k* = 0 to *k* = *k*_max_, we performed differentiation by annealing. If the outcome terminal cell corresponded to row *i* of Ξ, we adjusted *w*_*i*_ = *w*_*i*_ − 0.5(1 − *k/k*_max_), and then mean-adjusted **w** to zero.

## References

1. Burke, A. C., Nelson, C. E., Morgan, B. A., and Tabin, C. (1995). Hox genes and the evolution of vertebrate axial morphology. Development 121, 333–346.

2. Heinz, S., Benner, C., Spann, N., Bertolino, E., Lin, Y. C., Laslo, P., Cheng, J. X., Murre, C., Singh, H., and Glass, C. K. (2010). Simple combinations of lineage-determining transcription factors prime cis-regulatory elements required for macrophage and B cell identities. Molecular cell 38, 576–589.

3. Holmberg, J. and Perlmann, T. (2012). Maintaining differentiated cellular identity. Nature Reviews Genetics 13, 429–439.

4. Hnisz, D., Abraham, B. J., Lee, T. I., Lau, A., Saint-André, V., Sigova, A. A., Hoke, H. A., and Young, R. A. (2013). Super-enhancers in the control of cell identity and disease. Cell 155, 934–947.

5. Whyte, W. A., Orlando, D. A., Hnisz, D., Abraham, B. J., Lin, C. Y., Kagey, M. H., Rahl, P. B., Lee, T. I., and Young, R. A. (2013). Master transcription factors and mediator establish super-enhancers at key cell identity genes. Cell 153, 307–319.

6. Kelaini, S., Cochrane, A., and Margariti, A. (2014). Direct reprogramming of adult cells: avoiding the pluripotent state. Stem cells and cloning: advances and applications, 19–29.

7. Saint-André, V., Federation, A. J., Lin, C. Y., Abraham, B. J., Reddy, J., Lee, T. I., Bradner, J. E., and Young, R. A. (2016). Models of human core transcriptional regulatory circuitries. Genome research 26, 385–396.

8. Reiter, F., Wienerroither, S., and Stark, A. (2017). Combinatorial function of transcription factors and cofactors. Current opinion in genetics & development 43, 73–81.

9. Reilly, M. B., Cros, C., Varol, E., Yemini, E., and Hobert, O. (2020). Unique homeobox codes delineate all the neuron classes of C. elegans. Nature 584, 595–601.

10. Hobert, O. (2021). Homeobox genes and the specification of neuronal identity. Nature Reviews Neuroscience 22, 627–636.

11. Wang, H., Yang, Y., Liu, J., and Qian, L. (2021). Direct cell reprogramming: approaches, mechanisms and progress. Nature Reviews Molecular Cell Biology 22, 410–424.

12. Almeida, N., Chung, M. W., Drudi, E. M., Engquist, E. N., Hamrud, E., Isaacson, A., Tsang, V. S., Watt, F. M., and Spagnoli, F. M. (2021). Employing core regulatory circuits to define cell identity. The EMBO Journal 40, e106785.

13. Pennacchio, L. A., Bickmore, W., Dean, A., Nobrega, M. A., and Bejerano, G. (2013). Enhancers: five essential questions. Nature Reviews Genetics 14, 288–295.

14. Uyehara, C. M. and Apostolou, E. (2023). 3D enhancer-promoter interactions and multi-connected hubs: Organizational principles and functional roles. Cell Reports.

15. Calo, E. and Wysocka, J. (2013). Modification of enhancer chromatin: what, how, and why? Molecular cell 49, 825–837.

16. Park, Y.-K., Lee, J.-E., Yan, Z., McKernan, K., O’Haren, T., Wang, W., Peng, W., and Ge, K. (2021). Inter-play of BAF and MLL4 promotes cell type-specific enhancer activation. Nature communications 12, 1630.

17. Hansen, J. L., Loell, K. J., and Cohen, B. A. (2022). A test of the pioneer factor hypothesis using ectopic liver gene activation. Elife 11, e73358.

18. Heintzman, N. D., Hon, G. C., Hawkins, R. D., Kheradpour, P., Stark, A., Harp, L. F., Ye, Z., Lee, L. K., Stuart, R. K., Ching, C. W., et al. (2009). Histone modifications at human enhancers reflect global cell-type-specific gene expression. Nature 459, 108–112.

19. Creyghton, M. P., Cheng, A. W., Welstead, G. G., Kooistra, T., Carey, B. W., Steine, E. J., Hanna, J., Lodato, M. A., Frampton, G. M., Sharp, P. A., et al. (2010). Histone H3K27ac separates active from poised enhancers and predicts developmental state. Proceedings of the National Academy of Sciences 107, 21931– 21936.

20. Adam, R. C., Yang, H., Rockowitz, S., Larsen, S. B., Nikolova, M., Oristian, D. S., Polak, L., Kadaja, M., Asare, A., Zheng, D., et al. (2015). Pioneer factors govern super-enhancer dynamics in stem cell plasticity and lineage choice. Nature 521, 366–370.

21. Feng, C., Song, C., Jiang, Y., Zhao, J., Zhang, J., Wang, Y., Yin, M., Zhu, J., Ai, B., Wang, Q., et al. (2023). Landscape and significance of human super enhancer-driven core transcription regulatory circuitry. Molecular Therapy-Nucleic Acids 32, 385–401.

22. Heinz, S., Romanoski, C. E., Benner, C., and Glass, C. K. (2015). The selection and function of cell type-specific enhancers. Nature reviews Molecular cell biology 16, 144–154.

23. Sebé-Pedrós, A., Ballaré, C., Parra-Acero, H., Chiva, C., Tena, J. J., Sabidó, E., Gómez-Skarmeta, J. L., Di Croce, L., and Ruiz-Trillo, I. (2016). The dynamic regulatory genome of Capsaspora and the origin of animal multicellularity. Cell 165, 1224–1237.

24. Gaiti, F., Jindrich, K., Fernandez-Valverde, S. L., Roper, K. E., Degnan, B. M., and Tanurdžić, M. (2017). Landscape of histone modifications in a sponge reveals the origin of animal cis-regulatory complexity. Elife 6, e22194.

25. Ferrell Jr, J. E. (2002). Self-perpetuating states in signal transduction: positive feedback, double-negative feedback and bistability. Current opinion in cell biology 14, 140–148.

26. Huang, S., Guo, Y.-P., May, G., and Enver, T. (2007). Bifurcation dynamics in lineage-commitment in bipotent progenitor cells. Developmental biology 305, 695–713.

27. Alon, U. (2007). Network motifs: theory and experimental approaches. Nature Reviews Genetics 8, 450– 461.

28. Graham, T. G., Tabei, S. A., Dinner, A. R., and Rebay, I. (2010). Modeling bistable cell-fate choices in the Drosophila eye: qualitative and quantitative perspectives. Development 137, 2265–2278.

29. Sáez, M., Blassberg, R., Camacho-Aguilar, E., Siggia, E. D., Rand, D. A., and Briscoe, J. (2022). Statistically derived geometrical landscapes capture principles of decision-making dynamics during cell fate transitions. Cell Systems 13, 12–28.

30. Kauffman, S. (1969). Homeostasis and differentiation in random genetic control networks. Nature 224, 177– 178.

31. Huang, S., Ernberg, I., and Kauffman, S. (2009). “Cancer attractors: a systems view of tumors from a gene network dynamics and developmental perspective”. Seminars in cell & developmental biology. Vol. 20. Elsevier, 869–876.

32. Huang, S. (2012). The molecular and mathematical basis of Waddington’s epigenetic landscape: A frame-work for post-Darwinian biology? Bioessays 34, 149–157.

33. Lang, A. H., Li, H., Collins, J. J., and Mehta, P. (2014). Epigenetic landscapes explain partially reprogrammed cells and identify key reprogramming genes. PLoS computational biology 10, e1003734.

34. Fard, A. T., Srihari, S., Mar, J. C., and Ragan, M. A. (2016). Not just a colourful metaphor: modelling the landscape of cellular development using Hopfield networks. NPJ systems biology and applications 2, 1–9.

35. Zhu, R., Rio-Salgado, J. M. del, Garcia-Ojalvo, J., and Elowitz, M. B. (2022). Synthetic multistability in mammalian cells. Science 375, eabg9765.

36. Bintu, L., Buchler, N. E., Garcia, H. G., Gerland, U., Hwa, T., Kondev, J., and Phillips, R. (2005). Transcriptional regulation by the numbers: models. Current opinion in genetics & development 15, 116–124.

37. Tu, Y. (2013). Quantitative modeling of bacterial chemotaxis: signal amplification and accurate adaptation. Annual review of biophysics 42, 337–359.

38. Kvon, E. Z., Waymack, R., Gad, M., and Wunderlich, Z. (2021). Enhancer redundancy in development and disease. Nature Reviews Genetics 22, 324–336.

39. Hnisz, D., Schuijers, J., Lin, C. Y., Weintraub, A. S., Abraham, B. J., Lee, T. I., Bradner, J. E., and Young, R. A. (2015). Convergence of developmental and oncogenic signaling pathways at transcriptional super-enhancers. Molecular cell 58, 362–370.

40. Ramsauer, H., Schäfl, B., Lehner, J., Seidl, P., Widrich, M., Adler, T., Gruber, L., Holzleitner, M., Pavlović, M., Sandve, G. K., et al. (2020). Hopfield networks is all you need. arXiv preprint 2008.02217.

41. Kraut, R. and Levine, M. (1991). Mutually repressive interactions between the gap genes giant and Krüppel define middle body regions of the Drosophila embryo. Development 111, 611–621.

42. Milo, R., Shen-Orr, S., Itzkovitz, S., Kashtan, N., Chklovskii, D., and Alon, U. (2002). Network motifs: simple building blocks of complex networks. Science 298, 824–827.

43. Schaum, N., Karkanias, J., Neff, N. F., May, A. P., Quake, S. R., Wyss-Coray, T., Darmanis, S., Batson, J., Botvinnik, O., Chen, M. B., et al. (2018). Single-cell transcriptomics of 20 mouse organs creates a Tabula Muris: The Tabula Muris Consortium. Nature 562, 367.

44. D’Alessio, A. C., Fan, Z. P., Wert, K. J., Baranov, P., Cohen, M. A., Saini, J. S., Cohick, E., Charniga, C., Dadon, D., Hannett, N. M., et al. (2015). A systematic approach to identify candidate transcription factors that control cell identity. Stem cell reports 5, 763–775.

45. Rackham, O. J., Firas, J., Fang, H., Oates, M. E., Holmes, M. L., Knaupp, A. S., Consortium, F., Suzuki, H., Nefzger, C. M., Daub, C. O., et al. (2016). A predictive computational framework for direct reprogramming between human cell types. Nature genetics 48, 331–335.

46. Chanda, S., Ang, C. E., Davila, J., Pak, C., Mall, M., Lee, Q. Y., Ahlenius, H., Jung, S. W., Südhof, T. C., and Wernig, M. (2014). Generation of induced neuronal cells by the single reprogramming factor ASCL1. Stem cell reports 3, 282–296.

47. Merrell, A. J. and Stanger, B. Z. (2016). Adult cell plasticity in vivo: de-differentiation and transdifferentiation are back in style. Nature reviews Molecular cell biology 17, 413–425.

48. Moris, N., Pina, C., and Arias, A. M. (2016). Transition states and cell fate decisions in epigenetic landscapes. Nature Reviews Genetics 17, 693–703.

49. Månsson, R., Hultquist, A., Luc, S., Yang, L., Anderson, K., Kharazi, S., Al-Hashmi, S., Liuba, K., Thorén, L., Adolfsson, J., et al. (2007). Molecular evidence for hierarchical transcriptional lineage priming in fetal and adult stem cells and multipotent progenitors. Immunity 26, 407–419.

50. Orkin, S. H. and Zon, L. I. (2008). Hematopoiesis: an evolving paradigm for stem cell biology. Cell 132, 631– 644.

51. Kim, T.-H., Saadatpour, A., Guo, G., Saxena, M., Cavazza, A., Desai, N., Jadhav, U., Jiang, L., Rivera, M. N., Orkin, S. H., et al. (2016). Single-cell transcript profiles reveal multilineage priming in early progenitors derived from Lgr5+ intestinal stem cells. Cell reports 16, 2053–2060.

52. Qiao, X. T., Ziel, J. W., McKimpson, W., Madison, B. B., Todisco, A., Merchant, J. L., Samuelson, L. C., and Gumucio, D. L. (2007). Prospective identification of a multilineage progenitor in murine stomach epithelium. Gastroenterology 133, 1989–1998.

53. Toma, J. G., McKenzie, I. A., Bagli, D., and Miller, F. D. (2005). Isolation and characterization of multipotent skin-derived precursors from human skin. Stem cells 23, 727–737.

54. Briggs, J. A., Weinreb, C., Wagner, D. E., Megason, S., Peshkin, L., Kirschner, M. W., and Klein, A. M. (2018). The dynamics of gene expression in vertebrate embryogenesis at single-cell resolution. Science 360, eaar5780.

55. Waddington, C. H. (2014). The strategy of the genes. Routledge.

56. Hu, M., Krause, D., Greaves, M., Sharkis, S., Dexter, M., Heyworth, C., and Enver, T. (1997). Multilineage gene expression precedes commitment in the hemopoietic system. Genes & development 11, 774–785.

57. Miyamoto, T., Iwasaki, H., Reizis, B., Ye, M., Graf, T., Weissman, I. L., and Akashi, K. (2002). Myeloid or lymphoid promiscuity as a critical step in hematopoietic lineage commitment. Developmental cell 3, 137– 147.

58. Mercer, E. M., Lin, Y. C., Benner, C., Jhunjhunwala, S., Dutkowski, J., Flores, M., Sigvardsson, M., Ideker, T., Glass, C. K., and Murre, C. (2011). Multilineage priming of enhancer repertoires precedes commitment to the B and myeloid cell lineages in hematopoietic progenitors. Immunity 35, 413–425.

59. Olsson, A., Venkatasubramanian, M., Chaudhri, V. K., Aronow, B. J., Salomonis, N., Singh, H., and Grimes, H. L. (2016). Single-cell analysis of mixed-lineage states leading to a binary cell fate choice. Nature 537, 698–702.

60. Martin, E. W., Krietsch, J., Reggiardo, R. E., Sousae, R., Kim, D. H., and Forsberg, E. C. (2021). Chromatin accessibility maps provide evidence of multilineage gene priming in hematopoietic stem cells. Epigenetics & Chromatin 14, 1–15.

61. Zhou, J. X., Aliyu, M., Aurell, E., and Huang, S. (2012). Quasi-potential landscape in complex multi-stable systems. Journal of the Royal Society Interface 9, 3539–3553.

62. Rand, D. A., Raju, A., Sáez, M., Corson, F., and Siggia, E. D. (2021). Geometry of gene regulatory dynamics. Proceedings of the National Academy of Sciences 118, e2109729118.

63. Gulati, G. S., Sikandar, S. S., Wesche, D. J., Manjunath, A., Bharadwaj, A., Berger, M. J., Ilagan, F., Kuo, A. H., Hsieh, R. W., Cai, S., et al. (2020). Single-cell transcriptional diversity is a hallmark of developmental potential. Science 367, 405–411.

64. Choi, J., Baldwin, T. M., Wong, M., Bolden, J. E., Fairfax, K. A., Lucas, E. C., Cole, R., Biben, C., Morgan, C., Ramsay, K. A., et al. (2019). Haemopedia RNA-seq: a database of gene expression during haematopoiesis in mice and humans. Nucleic acids research 47, D780–D785.

65. Anderson III, D. A., Dutertre, C.-A., Ginhoux, F., and Murphy, K. M. (2021). Genetic models of human and mouse dendritic cell development and function. Nature Reviews Immunology 21, 101–115.

66. Arendt, D., Musser, J. M., Baker, C. V., Bergman, A., Cepko, C., Erwin, D. H., Pavlicev, M., Schlosser, G., Widder, S., Laubichler, M. D., et al. (2016). The origin and evolution of cell types. Nature Reviews Genetics 17, 744–757.

67. Hong, J.-W., Hendrix, D. A., and Levine, M. S. (2008). Shadow enhancers as a source of evolutionary novelty. Science 321, 1314–1314.

68. Cannavò, E., Khoueiry, P., Garfield, D. A., Geeleher, P., Zichner, T., Gustafson, E. H., Ciglar, L., Korbel, J. O., and Furlong, E. E. (2016). Shadow enhancers are pervasive features of developmental regulatory networks. Current Biology 26, 38–51.

69. Krotov, D. and Hopfield, J. (2020). Large associative memory problem in neurobiology and machine learning. arXiv preprint 2008.06996.

70. Amari, S.-I. (1972). Learning patterns and pattern sequences by self-organizing nets of threshold elements. IEEE Transactions on computers 100, 1197–1206.

71. Little, W. A. (1974). The existence of persistent states in the brain. Mathematical biosciences 19, 101–120.

72. Hopfield, J. J. (1982). Neural networks and physical systems with emergent collective computational abilities. Proceedings of the national academy of sciences 79, 2554–2558.

73. Guo, J. and Zheng, J. (2017). HopLand: single-cell pseudotime recovery using continuous Hopfield networkbased modeling of Waddington’s epigenetic landscape. Bioinformatics 33, i102–i109.

74. Conforte, A. J., Alves, L., Coelho, F. C., Carels, N., and Silva, F. A. B. d. (2020). Modeling basins of attraction for breast cancer using Hopfield networks. Frontiers in Genetics 11, 314.

75. Krotov, D. and Hopfield, J. J. (2016). Dense associative memory for pattern recognition. Advances in neural information processing systems 29.

76. Kirkpatrick, S., Gelatt Jr, C. D., and Vecchi, M. P. (1983). Optimization by simulated annealing. Science 220, 671–680.

77. Van Laarhoven, P. J., Aarts, E. H., Laarhoven, P. J. van, and Aarts, E. H. (1987). Simulated annealing. Springer.

78. Gao, H., Lukin, K., Ramírez, J., Fields, S., Lopez, D., and Hagman, J. (2009). Opposing effects of SWI/SNF and Mi-2/NuRD chromatin remodeling complexes on epigenetic reprogramming by EBF and Pax5. Proceedings of the National Academy of Sciences 106, 11258–11263.

79. Whyte, W. A., Bilodeau, S., Orlando, D. A., Hoke, H. A., Frampton, G. M., Foster, C. T., Cowley, S. M., and Young, R. A. (2012). Enhancer decommissioning by LSD1 during embryonic stem cell differentiation. Nature 482, 221–225.

80. Iurlaro, M., Stadler, M. B., Masoni, F., Jagani, Z., Galli, G. G., and Schübeler, D. (2021). Mammalian SWI/SNF continuously restores local accessibility to chromatin. Nature genetics 53, 279–287.

81. Wolf, B. K., Zhao, Y., McCray, A., Hawk, W. H., Deary, L. T., Sugiarto, N. W., LaCroix, I. S., Gerber, S. A., Cheng, C., and Wang, X. (2023). Cooperation of chromatin remodeling SWI/SNF complex and pioneer factor AP-1 shapes 3D enhancer landscapes. Nature Structural & Molecular Biology 30, 10–21.

82. Chan, H. L., Beckedorff, F., Zhang, Y., Garcia-Huidobro, J., Jiang, H., Colaprico, A., Bilbao, D., Figueroa, M. E., LaCava, J., Shiekhattar, R., et al. (2018). Polycomb complexes associate with enhancers and promote oncogenic transcriptional programs in cancer through multiple mechanisms. Nature communications 9, 3377.

83. Angeloni, A. and Bogdanovic, O. (2019). Enhancer DNA methylation: implications for gene regulation. Essays in biochemistry 63, 707–715.

84. Blomen, V. and Boonstra, J. (2007). Cell fate determination during G1 phase progression. Cellular and Molecular Life Sciences 64, 3084–3104.

85. Dalton, S. (2015). Linking the cell cycle to cell fate decisions. Trends in cell biology 25, 592–600.

86. Cermakova, K., Tao, L., Dejmek, M., Sala, M., Montierth, M. D., Chan, Y. S., Patel, I., Chambers, C., Loeza Cabrera, M., Hoffman, D., et al. (2023). Reactivation of the G1 enhancer landscape underlies core circuitry addiction to SWI/SNF. Nucleic Acids Research, gkad1081.

